# The impact of genistein supplementation on tendon functional properties and gene expression in estrogen deficient rats

**DOI:** 10.1101/715102

**Authors:** Chad C. Carroll, Shivam H. Patel, Jessica Simmons, Ben DH. Gordon, Jay F. Olson, Kali Chemelewski, Shannon Saw, Taben M. Hale, Reuben Howden, Arman Sabbaghi

## Abstract

**Purpose:** Tendinopathy risk increases with menopause. The phytoestrogen genistein prevents collagen loss during estrogen deficiency [ovariectomy (OVX)]. The influence of genistein on tendon function and extracellular matrix (ECM) regulation are not well known. We determined the impact of genistein on tendon function and examined potential mechanisms by which genistein alters tendon ECM.

**Materials and Methods:** Eight-week-old rats (n=42) were divided into three groups: intact, OVX, or OVX-genistein (6mg/kg/day) for 6-weeks. Tail fascicles were assessed with a Deben tensile stage. Achilles tendon mRNA expression was determined with digital droplet PCR. Tendon-derived fibroblasts were also treated with genistein in the presence of estrogen receptor (ER) antagonists.

**Results:** Compared to intact, stress tended to be lower in untreated OVX rats (p=0.022). Further, modulus and energy density were greater in genistein-treated rats (p<0.05) compared to intact. Neither OVX nor genistein altered expression of Col1a1, Col3a1, Casp3, Casp8, Mmp1a, Mmp2, or Mmp9 (p>0.05). Compared to intact, Tnmd and Esr1 expression was greater and Pcna and Timp1 expression lower in OVX rats (p<0.05). Genistein treatment returned Tnmd, Pcna, and Timp1 to levels of Intact-Vehicle (p<0.05), but did not alter Scx or Esr1 (p>0.05). Several β-catenin/Wnt signaling related molecules were not altered by OVX or genistein (p>0.05). *In vitro*, genistein blunted cell proliferation but not via ERs.

**Conclusions:** Our findings demonstrate that genistein improves tendon function. Genistein inhibits cell proliferation *in vitro* but not via ER. The effect of genistein *in vivo* was predominately on genes related to cell proliferation rather than collagen remodeling.

## INTRODUCTION

The transition through menopause is a significant risk factor for tendinopathies (tendon pain/degeneration), an understudied clinical problem (Abate et al. 2014). Menopause also leads to deteriorations in tendon function (Frizziero et al. 2014), changes that likely contribute to the global decline in musculoskeletal function associated with menopause (Nedergaard et al. 2013). Overall, tendon alterations during menopause contribute to significant physical impairment, reduced quality of life, and high socio-economic costs (Hopkins et al. 2016). Although it is unclear if the loss of estrogen with menopause is the primary reason for alterations in tendon properties, estrogen appears to be vital for maintenance of tendon extracellular matrix [ECM; (Cook et al. 2007; Hansen et al. 2008; Hansen, Kongsgaard, et al. 2009; Hansen, Miller, et al. 2009; Ramos et al. 2012)]. Specifically, we have shown that in a rat model of estrogen deficiency [ovariectomy (OVX)], estrogen loss results in a large decline in tendon collagen content (Ramos et al. 2012), the primary structural protein of tendon. Further, in tendon cell culture, estrogen deficiency results in decreased cell proliferation, reduced type I collagen (Torricelli et al. 2013), and altered tendon metabolism and healing more than attributable to aging alone (Torricelli et al. 2013). Estrogen loss may also lead to increased activity of ECM degrading enzymes (Pereira et al. 2010). Tissue collagen content directly correlates with tissue stiffness (Ducomps et al. 2003), implying that a reduction in tendon collagen could contribute to a reduction in tendon functional properties. However, the impact of estrogen loss on tendon functional properties has not been extensively studied (Circi et al. 2009). Further, the molecular mechanisms modulating the effects of estrogen on tendon are not well defined.

A common therapeutic choice for estrogen-deficiency, primarily postmenopause, is estrogen therapy (ET). However, in postmenopausal women ET is associated with tendons that have a lower Young’s modulus and a greater number of smaller collagen fibrils (Hansen, Kongsgaard, et al. 2009). ET also limits tendon hypertrophy with exercise in postmenopausal women (Cook et al. 2007; Finni et al. 2009). Moreover, it has been suggested that tendinopathy is more common with ET compared to no ET (Holmes and Lin 2006). Collectively, these findings suggest that alternative approaches to minimizing the impact of estrogen loss on tendon are needed.

Genistein, a plant-derived, naturally occurring isoflavone phytoestrogen found in soy, may be a useful strategy to reduce the effects of estrogen deficiency on tendons (Ramos et al. 2012). Genistein is thought to convey many of the health benefits associated with a diet high in soy, including improved bone mass (Marini et al. 2007; Qi and Zheng 2017), reduced breast cancer risk (Trock et al. 2006; Verheus et al. 2007), and reduced risk of cardiovascular disease (Squadrito et al. 2003). The beneficial effects of genistein on bone and cardiovascular disease remain controversial (Abdi et al. 2016), but in an OVX rat model we demonstrated that genistein reverses the decline in tendon collagen associated with the loss of estrogen (Ramos et al. 2012). However, the mechanisms by which genistein modulates ECM have not been explored and it is unclear if genistein can improve tendon mechanical properties in estrogen deficient rats. The objectives of this investigation were to: 1) determine if genistein can improve tendon functional properties in estrogen deficient rats, and 2) examine the impact of estrogen loss and genistein supplementation on key regulators of ECM that could account for the changes in tendon collagen noted in our previous work (Ramos et al. 2012).

## MATERIALS AND METHODS

### Study Protocol

Eight-week-old Sprague-Dawley rats were purchased from Charles River Laboratories (Wilmington, MA) as either OVX (n=28) or Intact (n=14) (Ramos et al. 2012). OVX rats were randomly assigned to one of two groups: OVX-Vehicle or OVX-Genistein. OVX rats received a daily subcutaneous injection of either six mg/kg/day of genistein (LC Laboratories, Woburn, MA, USA) in DMSO (OVX-Genistein) or DMSO only (OVX-Vehicle). Intact rats received a daily injection of DMSO (Intact-Vehicle) equivalent to the quantity of DMSO delivered to OVX rats. The chosen dose of genistein was designed to represent a human diet high in soy, which has been associated with lower rates of breast and prostate cancer (Tham et al. 1998) and modest changes in risk factors for cardiovascular disease and osteoporosis (Anderson et al. 1998) without effects on endometrial thickness (Alekel et al. 2014). Doses were scaled to a rat equivalent (Services 2005). Rats were studied in two separate cohorts with 18 rats (6 per group) in cohort 1 and 24 rats (8 per group) in cohort 2. Tail tendon fascicle mechanical properties were evaluated in all rats; however, tendons from cohort 1 were used for a separate set of RNA sequencing experiments. Thus, gene expression data is reported from eight rats per group.

After group assignment, rats were caged in pairs (with each pair being members of the same experimental group), allowed access to genistein-free food (Modified AIN-93G with Corn Oil, Dyets Inc, Bethlehem, PA), water ad libitum, and maintained on a 12-hour light-dark cycle. After a two-week equilibration period, rats were treated with genistein or vehicle daily for six weeks. After six weeks, rats were euthanized by decapitation after CO_2_ inhalation (Ramos et al. 2012). Tendons were carefully dissected from the rats, removed of any excess fascia and muscle, flash frozen in liquid nitrogen, and transferred to −80°C. Tails were immediately removed, wrapped in saline soaked gauze, and stored at −20°C until analysis. This investigation was approved by the Purdue University and Midwestern University Institutional Animal Care and Use Committees and all animals were cared for in accordance with the recommendations in the Guide for the Care and Use of Laboratory Animals (Council 2011).

### Achilles Tendon Gene Expression

Total RNA for gene expression analysis was isolated from the Achilles tendon as previously described (Patel et al. 2018). Briefly, 10 to 20 mg of frozen tendon tissue was pulverized under cryogenic conditions (OPS Diagnostics, Lebanon, NJ) and lysed using a bead mill homogenizer (BR12, Omni) in TRIzol Reagent (Invitrogen). Phase separation and RNA precipitation were completed per the manufacturer’s instructions (TRIzol, Invitrogen). RNA concentration was determined using a NanoDrop 2000 (Thermo Fisher Scientific). Quality of RNA was assessed using the 260/280 and 260/230 ratios. Reverse transcription (iScript, BioRad, Hercules, CA) was completed to produce complementary DNA from 100ng of RNA. Absolute quantification of mRNA target transcripts was completed using digital droplet PCR (ddPCR; BioRad) and reported as positive counts per 20µl reaction (Patel et al. 2018). Input cDNA for all genes was 1.11 ng, excluding Col1a1, Col3a1, Tnmd, and Dcn, which were 0.55 ng.

### Tail Tendon Fascicle Mechanical Properties

Tail fascicles (~30 mm in length) were harvested and their mechanical properties were assessed as previously described (Volper et al. 2015). Briefly, we used a Deben mini tensile tester 200 N stage (Deben, UK, Ltd.) and a Nikon SMZ1000 microscope with a color C-mos chip 1280×1024 Pixelink camera attached for sample imaging. Tensile force was applied by increasing the distance between the clamps at 1mm/min from resting (0.20 N) to ~0.25 N. Ten cycles for pre-conditioning were performed during which fascicles were subjected to 3% deformation from resting length. Following pre-conditioning, images were captured from the top and side for calculation of cross-sectional area and tendon fascicle length at the onset of force (0.01-0.05 N) with fascicle crimps evaluated to be intact. Sample mechanical properties were then determined by increasing the distance between the clamps at 6 mm/min until sample failure while recording force data at 10 Hz. Force and dimension data were subsequently used to calculate maximum force, maximum stress, and Young’s modulus, each of which were determined as the peak value of a moving linear regression over 10 data points.

### Histology

Immersion-fixed (Histo-choice Fixative D, Amresco) longitudinal sections of the Achilles tendon were dehydrated through graded methanol, cleared with toluene, and embedded in paraffin (Formula R, Leica Microsystems Surgipath, Buffalo Grove, IL) (Volper et al. 2015). Sections (4 μm) were dewaxed and rehydrated through graded ethanol and stained with hematoxylin and eosin (H&E) staining to determine cell density (Volper et al. 2015). Cell counting was completed independently by two investigators.

### Cell Culture

Tendon-derived fibroblasts were isolated from the Achilles tendon of four rats from the Intact group. After euthanasia, tendon was rinsed with sterile PBS, finely minced, placed in DMEM containing 0.2% type I collagenase, and incubated in a 37°C shaking water bath for four hours. After tissue digestion, the cell suspension was filtered through a 100µm mesh filter, pelleted by centrifugation, resuspended in 5.5mM glucose DMEM containing 10% FBS, 1% sodium pyruvate (Sigma), and 1% penicillin/streptomycin (Thermo Scientific) and plated in 100mm collagen coated dishes. After reaching ~75-80% confluence, tendon fibroblasts were seeded with media containing DMSO vehicle, genistein (150 μM, LC Laboratories), or estradiol (1.1 μM, Sigma-Aldrich) and were treated with control, MPP (100 μM, Sigma-Aldrich), or PHTPP (1000 μM, Sigma-Aldrich) for 48 hours. Tendon fibroblasts from passage 2-4 were used for all experiments. Cell counts were completed in duplicate using a hemocytometer.

### Statistical Analysis

Our statistical analyses on contrasts between the Intact-Vehicle, OVX-Vehicle, and OVX-Genistein groups proceed via two classes of regression models. Normal linear regression models were fit for the tendon gene expressions data, and mixed effects regression models were fit for the tail fascicle mechanical properties and body weight data. The independent variables that were considered for all of the regression models were indicators for the two OVX groups, with the intact group serving as the baseline. Further details on the specific regression analyses that were performed for these three types of dependent variables are provided in the following subsections.

#### Regression Analysis for Tendon Gene Expressions

Separate Normal linear regression models were fit for each individual gene using the lm function in the statistical software environment R. For each fitted model, we checked whether the standard assumptions for linear regression were satisfied on the original scale of the dependent variable. If so, we then performed tests for all pairwise contrasts between the three groups using the regression parameters’ estimators and standard errors. If the assumptions were not satisfied, we then considered a Normal linear regression model on the square root scale of the dependent variable (which is a standard variance-stabilizing transformation for the count data inherent in gene expressions), as well as weighted linear regression models on both the original and square root scales. We then determined which of these latter three models provided the best fit to the data in terms of satisfying the regression assumptions. Once a final model was selected, we proceeded to perform tests for all of the pairwise contrasts. Our chosen significance level was α = 0.05 throughout, and we accounted for the multiple comparisons among the three groups by means of separate Bonferroni adjustments across the genes.

#### Mixed Effects Regression Modeling for Mechanical Properties

Separate mixed effects regression models were fit for each mechanical property using the lmer function in the lme4 package in R. As the mechanical properties data were collected from two distinct cohorts of rats, each model had two sets of fixed effects that correspond to the OVX and cohort groups, and one set of random effects that correspond to unique, random intercepts for the rats. These intercepts are assumed to be independent Normal random variables with mean 0 and identical variance. This specification of a single model with fixed and random effects can effectively account for the expected correlation in the properties of the two fascicles that were extracted from one rat, as well as possible differences that arise across the cohorts.

The analysis for each property proceeds via consideration of two mixed effects regression models, referred to as the “main effects” and “full” models, respectively. Both have fixed effects for cohort, OVX-Vehicle, and OVX-Genistein, and random intercepts for the rats. However, the full model considers interactions between cohort and OVX, whereas the main effects model does not. We first tested the significance of the interactions in the full model. If any interaction is statistically significant we selected the full model, whereas if no interaction is statistically significant we selected the main effects model. In either case, we diagnosed the validity of the assumptions for the selected mixed effects regression model. If the assumptions were satisfied, we tested all three pairwise main effect contrasts for the treatments. If the assumptions were not satisfied, we performed the same type of mixed effects regression analyses described above on the logarithmic-transformed dependent variable. As before, our chosen significance level was α = 0.05 throughout, with a Bonferroni correction made to adjust for the fact that we are testing three contrasts.

#### Mixed Effects Longitudinal Regression Modeling for Body Weight

A mixed effects longitudinal regression model was fit for the rat body weight data using the lmer function in the lme4 package in R. The body weights were collected on the previously described two cohorts of rats. As body weight is a positive, continuous measurement, we fitted the mixed effects model on the logarithmic-transformed weights to better satisfy the regression assumptions. On the basis of exploratory data analyses and visualizations, we compared two mixed effects regression model specifications to determine how rat body weight depends on the different treatments and cohorts, and changes across time: a “full” model that incorporates fixed main effects and two-factor interactions for OVX groups, cohort, and time, and a “reduced” model that incorporates fixed main effects and two-factor interactions only for OVX groups and time. In both of these models, a random intercept and a random slope for time is introduced for each rat. These random effects are specified to capture observed variations in linear time trends and intercepts between rats. The validity of the assumptions for the mixed effects longitudinal regression models were assessed via residual diagnostics. After fitting these two mixed effects regression model, we tested the significance of the interactions and determined whether the reduced model was sufficient for explaining the observed trends in the body weight data. Finally, we tested the fixed effects for the selected regression model.

## RESULTS

Body mass increased with time in all groups but rats in the OVX groups were heavier than intact rats (p<0.05, Figure 1). Tendon tail fascicle stiffness was not altered by OVX or genistein treatment (p>0.05, Table 2). Deformation (p=0.048), maximum stress (p=0.022) and yield stress (p=0.052) tended to be lower in OVX-Vehicle rats when compared to Intact-Vehicle but failed to reach statistical significance after correction for multiple comparisons (adjusted α=0.017, Table 2). In contrast, energy density at yield, modulus, maximum stress, and yield stress were greater in OVX-Genistein rats when compared to OVX-Vehicle (p<0.05, Table 2).

**Table 1:** ddPCR Probes

| **Gene Symbol** | **Gene Name** | **Entrez Gene** |
| --- | --- | --- |
| Casp3 | Caspase-3 | 25402 |
| Casp8 | Caspase-8 | 64044 |
| Ccnd1 | G1/S-specific cyclin-D1 | 58919 |
| Col1a1 | Collagen alpha-1(I) chain | 29393 |
| Col3a1 | Collagen alpha-1(III) chain | 84032 |
| Ctnnb1 | Catenin beta-1 | 84353 |
| Dcn | Decorin | 29139 |
| Dkk1 | Dickkopf-like protein 1 | 22943 |
| Esr1 | Estrogen receptor $\alpha$ | 24890 |
| Esr2 | Estrogen receptor $\beta$ | 25149 |
| Mmp1a | Matrix metalloproteinase 1a precursor | 300339 |
| Mmp2 | Matrix metalloproteinase 2 | 81686 |
| Mmp9 | Matrix metalloproteinase 9 | 81687 |
| Pcna | Proliferating cell nuclear antigen | 25737 |
| Scx | Scleraxis | 680712 |
| Tgfbr3 | Transforming growth factor beta receptor type 3 (betaglycan) | 29610 |
| Timp1 | Tissue inhibitor of matrix metalloproteinase 1 | 116510 |
| Tnmd | Tenomodulin | 64104 |
| Wnt16 | Protein Wnt16 precursor | 500047 |

**Table 2:** Tail Tendon Fascicle Mechanical Properties

|  | **Intact-Vehicle** | **OVX-Vehicle** | **OVX-Genistein** |
| --- | --- | --- | --- |
| **Deformation (mm)** | 2.62±0.59 | 2.32±0.50 | 2.37±0.46 |
| **Energy Density at Yield (MJ/m<sup>3</sup>)** | 1.56±0.63 | 1.36±0.46 | 1.72±0.36* |
| **Stiffness (N/mm)** | 1.33±0.64 | 1.38±0.61 | 1.49±0.66 |
| **Young's modulus (MPa)** | 1169±285 | 1074±277 | 1244±240* |
| **Maximum Stress (MPa)** | 60.5±17.7 | 52.1±12.9 | 62.1±12.3* |
| **Yield Stress (MPa)** | 49.8±16.2 | 43.2±13.7 | 53.9±9.8* |
| **Strain (%)** | 0.061±0.009 | 0.058±0.011 | 0.062±0.005 |

Data are presented as means ± SE; *n* = 14 rats per group.
\**p*<0.05 vs. OVX-Vehicle

**Figure 1:**
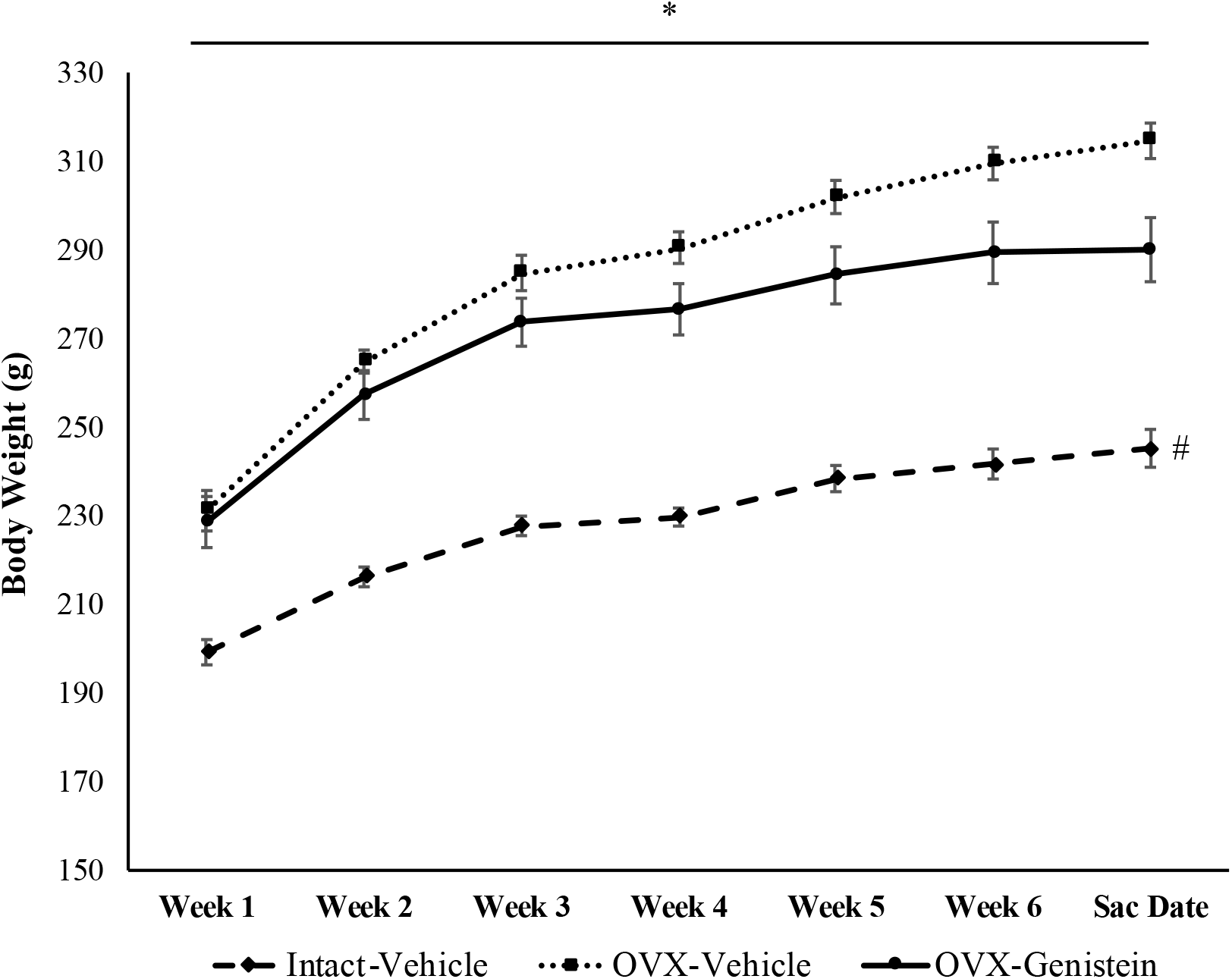
Rat body mass (g) over the course of the treatment protocol (n=14). Rats were weighed weekly. *p<0.05 main effect for time, all groups. ^#^p<0.05, Intact-Vehicle lower than OVX-Vehicle and OVX-Genistein at all time points. Data presented as mean and standard error

Table 3 provides list of the genes evaluated ranked by total counts in the Achilles tendon of Intact rats, providing an indication of the relative abundance of each gene transcript evaluated. Dcn and Col1a1 were the genes demonstrating the greatest expression levels. Neither Col1a1 (Intact-Vehicle: 85045±20981 counts per 20 μl; OVX-Vehicle: 95225±7549; OVX-Genistein: 71380±13846) nor Col3a1 (Intact-Vehicle: 10568±1007; OVX-Vehicle: 10243±1112; OVX-Genistein: 9070±810) expression was affected by OVX or genistein treatment (Figure 2). Decorin was not influenced by OVX (Intact-Vehicle: 96600±6272; OVX-Vehicle: 80625±9301) but was 36% greater in genistein-treated rats (OVX-Genistein: 115800±4222) when compared to OVX-Vehicle (p<0.05, Figure 2). In contrast, Tgfbr3 (betaglycan) was not influenced by OVX or genistein (Intact-Vehicle: 1889±403; OVX-Vehicle: 1432±198; OVX-Genistein: 1521±225).

**Table 3:**
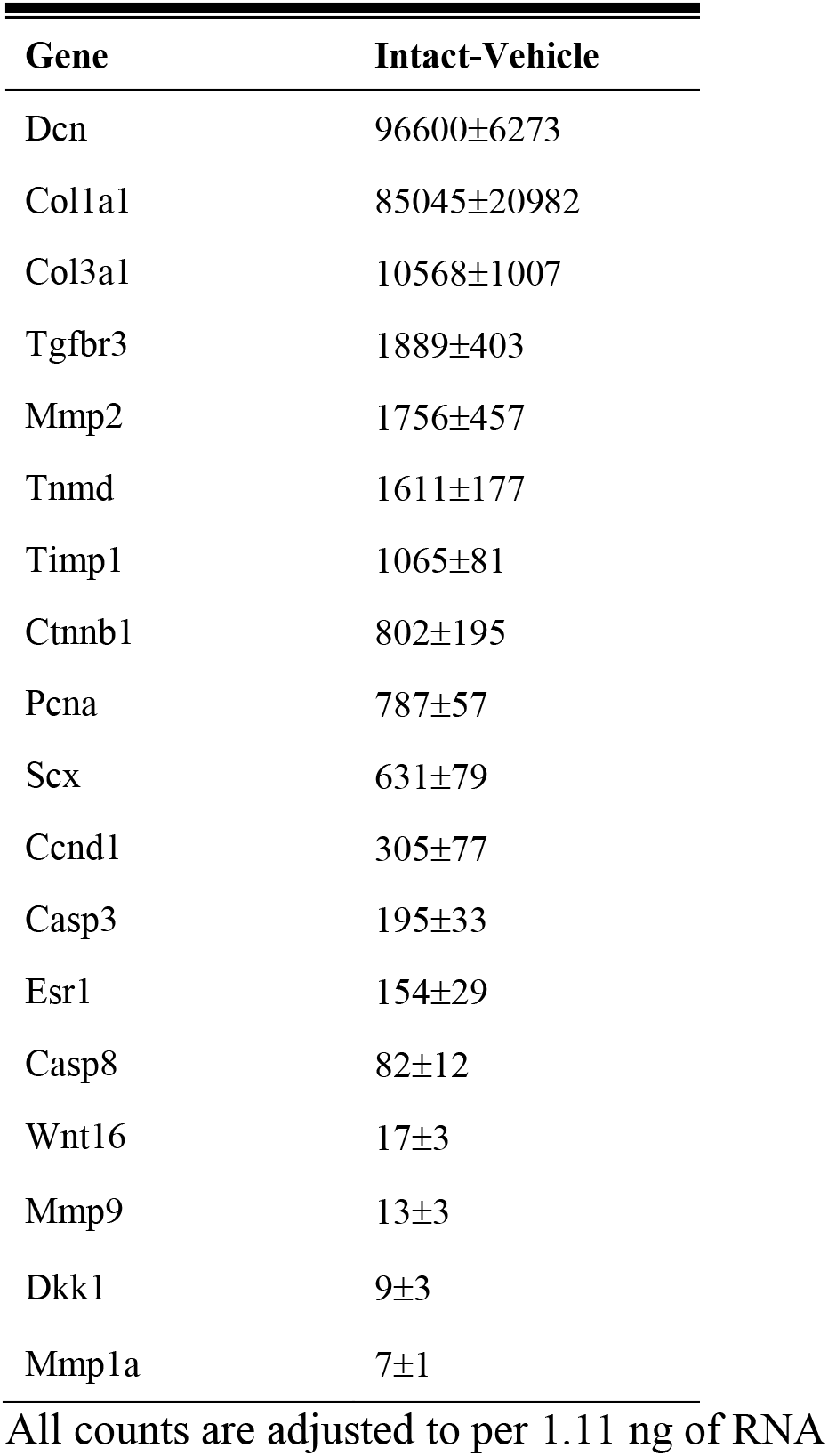
Genes Ranked by Counts in Intact-Vehicle Rats

**Figure 2:**
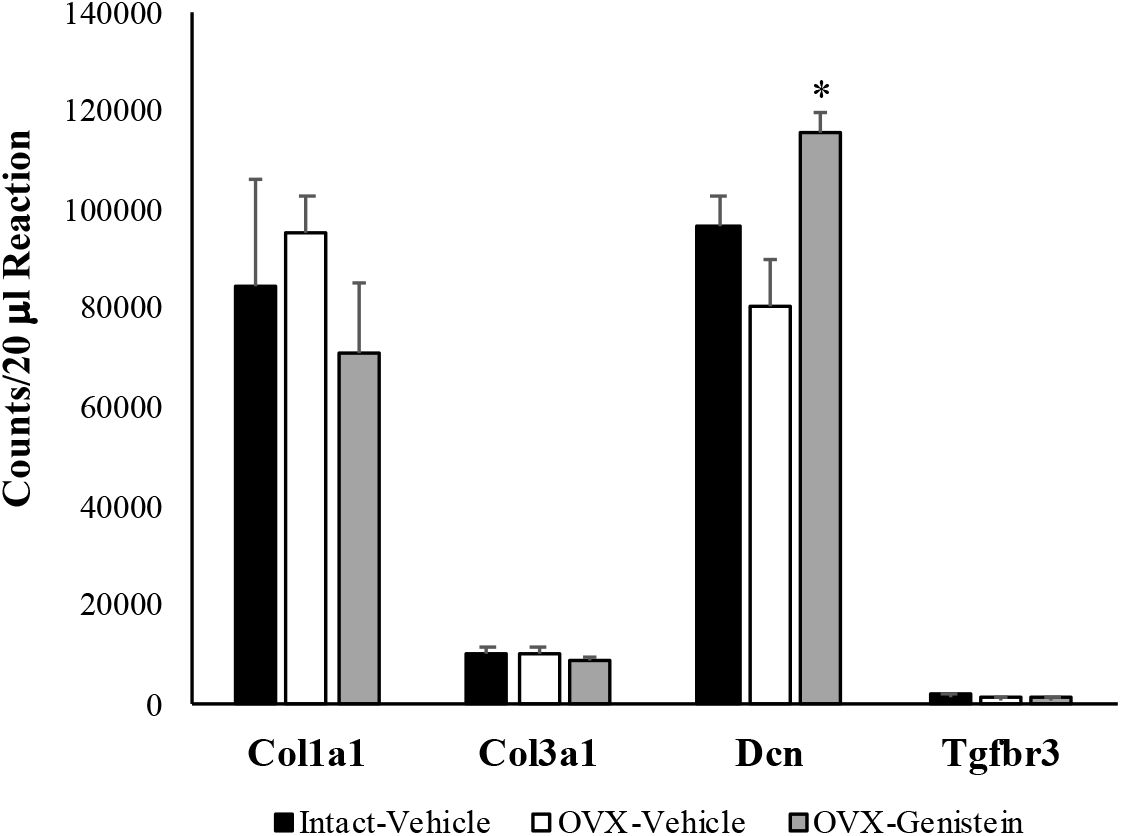
ddPCR counts for collagen 1 (Col1a1), collagen 3 (Col3a1), decorin (Dcn), and transforming growth factor beta receptor type 3 or betaglycan (Tgfbr3). *p<0.05 vs. OVX-Vehicle. Data presented as mean and standard error for transcript counts per 20 μl reaction.

Gene expression of the ECM enzymes Mmp1a (Intact-Vehicle: 7±1; OVX-Vehicle: 10±3; OVX-Genistein: 12±2), Mmp2 (Intact-Vehicle: 1756±457; OVX-Vehicle: 2369±370; OVX-Genistein: 1749±326), Mmp9 (Intact-Vehicle: 13±3; OVX-Vehicle: 12±3; OVX-Genistein: 15±2), and Timp1 (Intact-Vehicle: 1065±81; OVX-Vehicle: 939±95; OVX-Genistein: 1318±82) were not influenced by OVX or genistein treatment (p>0.05, Figure 3). In contrast, Tnmd expression was 2.4-fold greater (p<0.05, Figure 3) in OVX-Vehicle (3943±442) when compared to Intact-Vehicle (1611±177). Tnmd expression was 45% lower in OVX-Genistein (2498±115) rats when compared to OVX-Vehicle, but still higher than Intact-Vehicle (p<0.05, Figure 4). Further, Pcna expression was 33% lower in OVX-Vehicle (563±20) in comparison to Intact-Vehicle (787±57, p<0.05) whereas the reduction was prevented by genistein treatment (Figure 4, 734±48). Scx expression was not altered by OVX or genistein treatment (p>0.05, Figure 4, Intact-Vehicle: 631±79; OVX-Vehicle: 961±85; OVX-Genistein: 1001±145).

**Figure 3:**
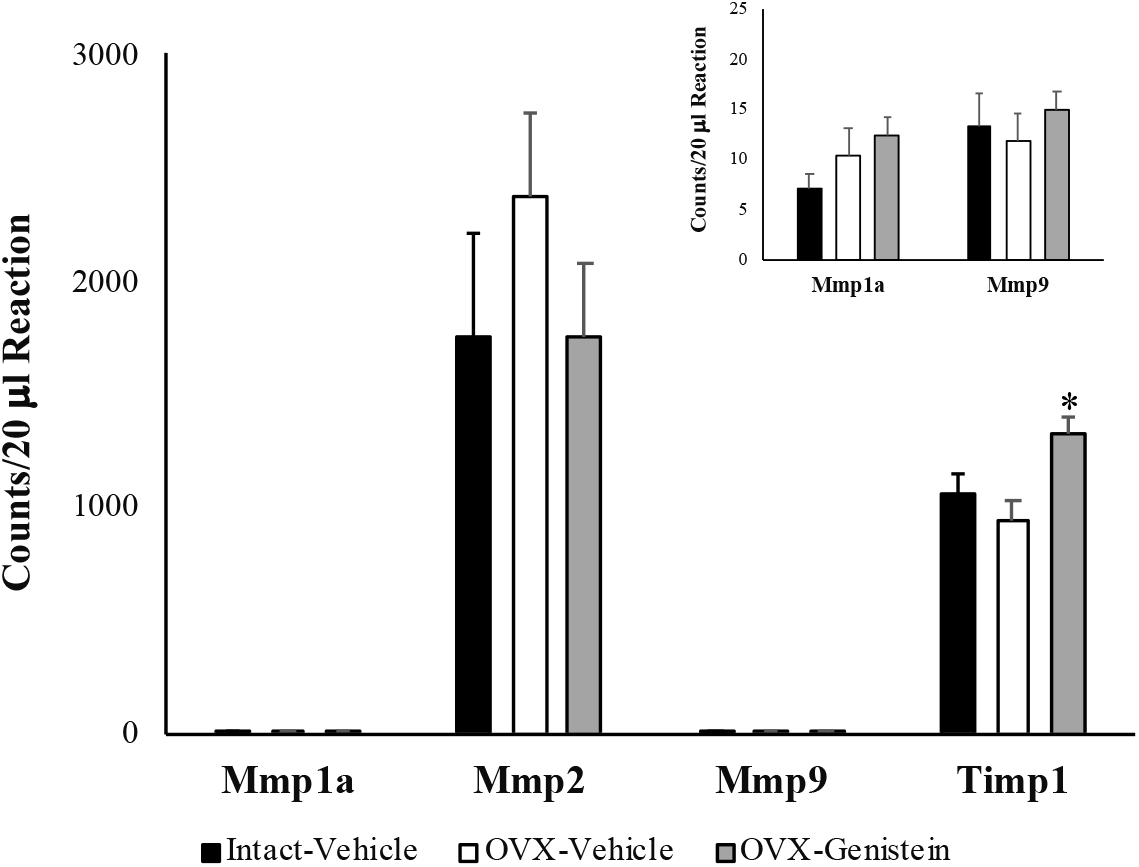
ddPCR counts for matrix metalloproteinase 1, 2, and 9 (Mmp), tissue inhibitor of MMP 1 (Timp1), *p<0.05 vs. OVX-Vehicle. Data presented as mean and standard error for transcript counts per 20 μl reaction. Due to the low counts of Mmp 1a and Mmp9, data is also shown to a smaller scale in upper right of figure.

**Figure 4:**
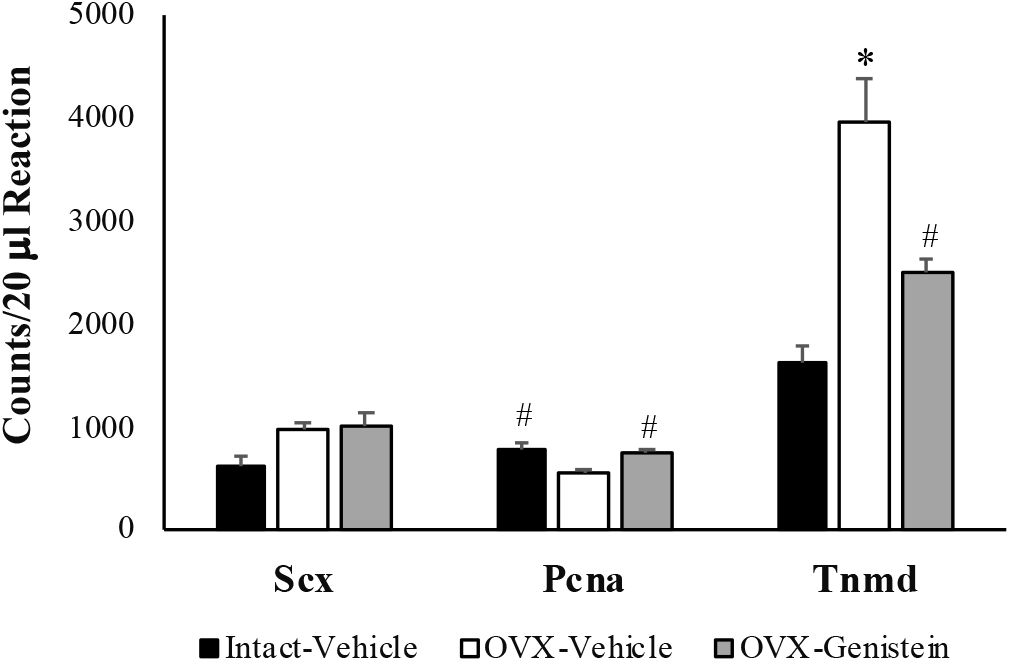
ddPCR counts for tenomodulin (Tnmd), scleraxis (Scx), and proliferating cell nuclear antigen (Pcna). *p<0.05 vs. OVX-Vehicle and OVX-Genistein. ^#^p<0.05 vs. OVX-Vehicle. Data presented as mean and standard error for transcript counts per 20 μl reaction.

The apoptotic genes Casp3 (Intact-Vehicle: 195±33; OVX-Vehicle: 244±11; OVX-Genistein: 221±21) and Casp8 (Intact-Vehicle: 82±12; OVX-Vehicle: 91±13; OVX-Genistein: 100±10) were not influenced by OVX or genistein (p>0.05, Figure 5). Esr1 expression was greater (p<0.05, Figure 5) in OVX-Vehicle (307±47) compared to Intact-Vehicle (154±29). Esr1 expression also tended to be higher (2-fold) in OVX-Genistein rats (271±31, p=0.023) when compared to Intact-Vehicle, but this effect did not reach statistical significance after correcting for the multiple comparisons. Esr2 transcripts were not detected using the predesigned Biorad probe. Wnt16 (Intact-Vehicle: 17±3; OVX-Vehicle: 34±6; OVX-Genistein: 26±5), Ccnd1 (Intact-Vehicle: 305±77; OVX-Vehicle: 385±47; OVX-Genistein: 420±88), Ctnnb1 (Intact-Vehicle: 802±195; OVX-Vehicle: 966±134; OVX-Genistein: 875±160), and Dkk1 (Intact-Vehicle: 9±3; OVX-Vehicle: 10±2; OVX-Genistein: 9±3) expression were not altered by OVX or genistein treatments (p>0.05, Figure 6).

**Figure 5:**
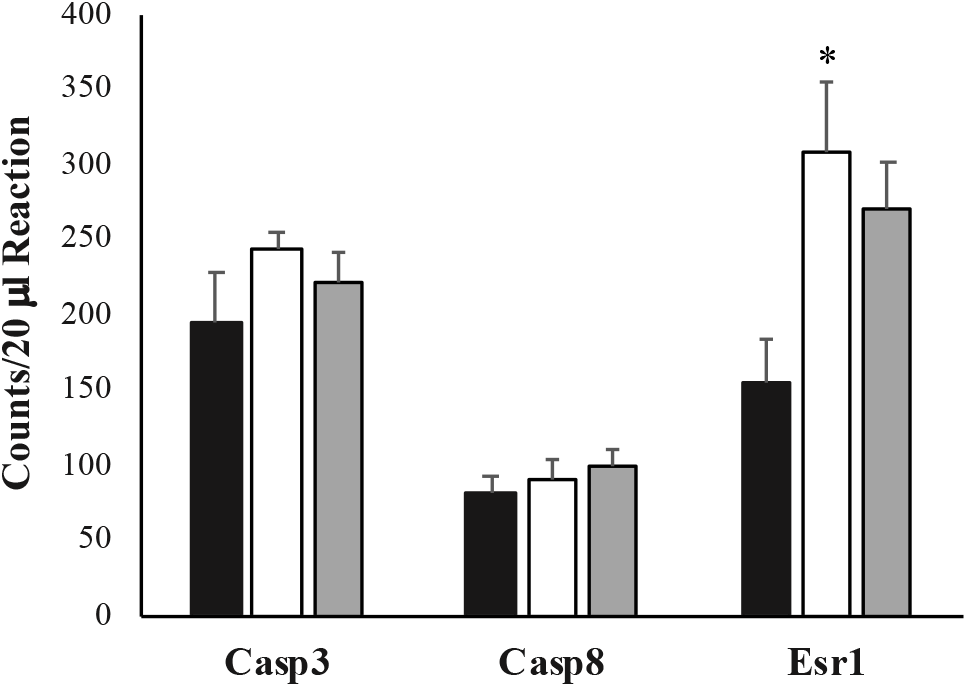
ddPCR counts for caspase 3 and 9 (Casp) and estrogen receptor 1 (Esr1). *p<0.05 vs. OVX-Vehicle. Data presented as mean and standard error for transcript counts per 20 μl reaction.

**Figure 6:**
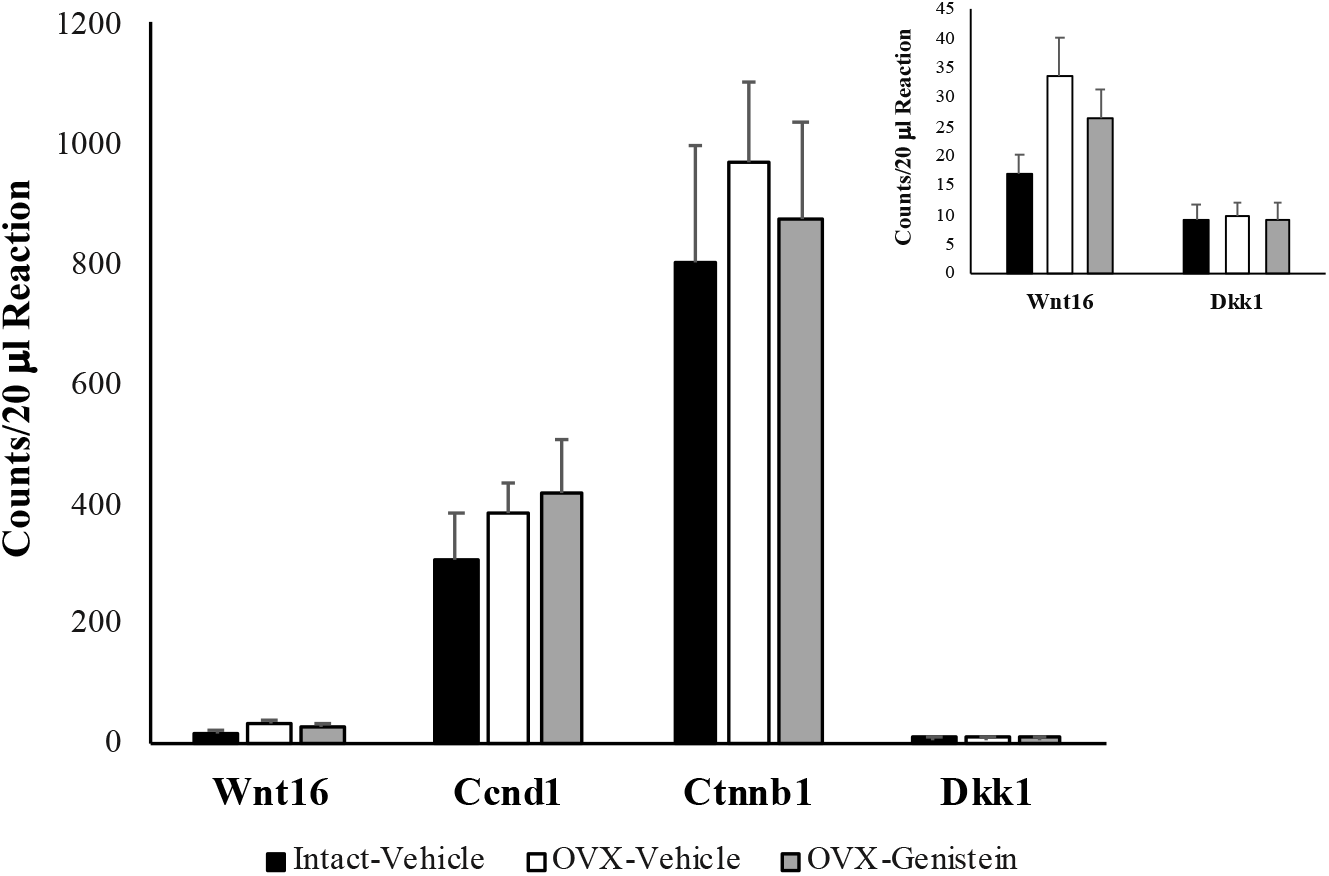
ddPCR counts for Wnt family member 16 (Wnt16), cyclin D1 (Ccnd1), catenin beta 1 (Ctnnb1), and Dickkopf WNT Signaling Pathway Inhibitor 1 (Dkk1). Data presented as mean and standard error for transcript counts per 20 μl reaction. Due to the low counts of Wnt16 and Dkk1, data is also shown to a smaller scale in upper right of figure.

**Figure 7:**
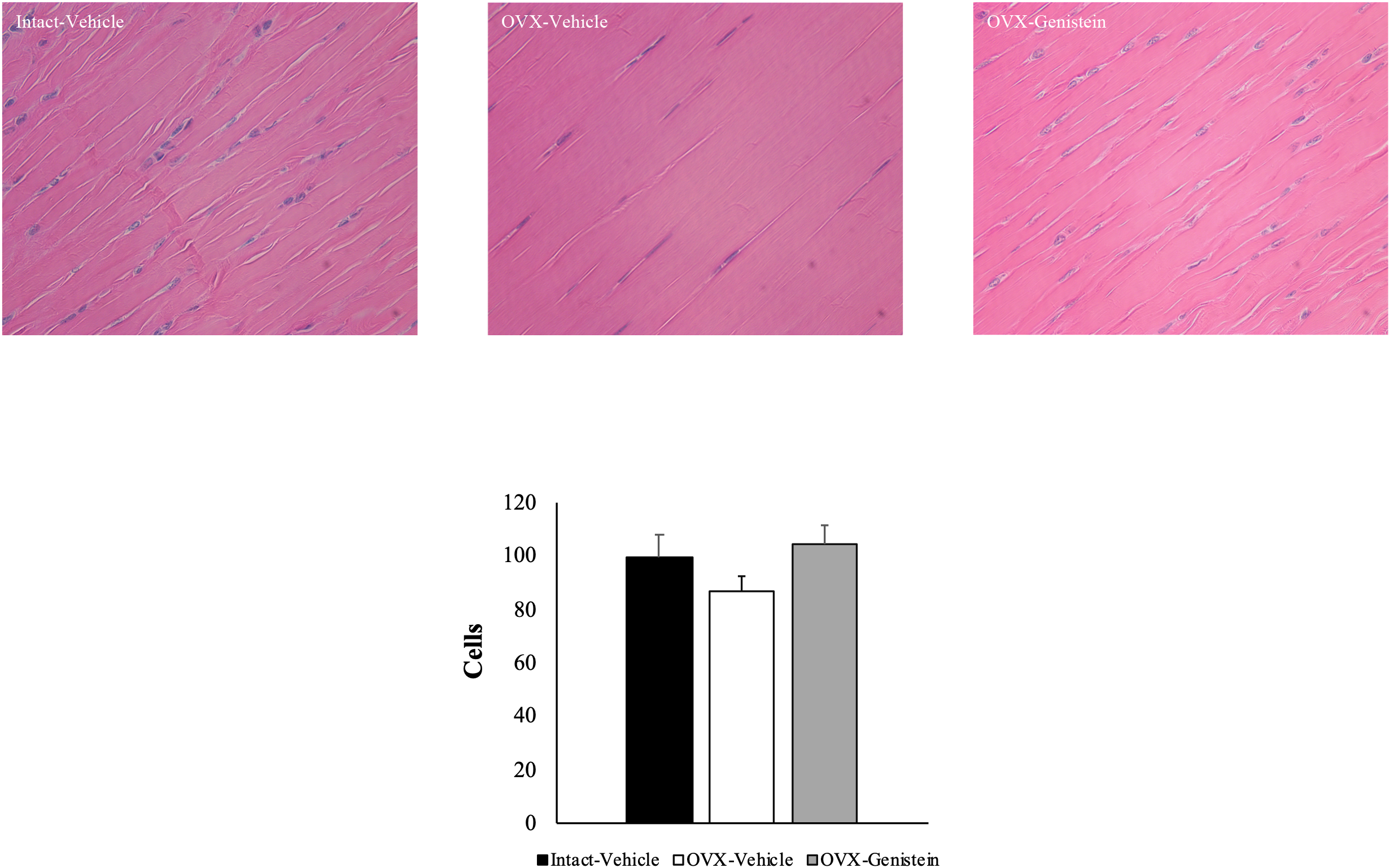
Achilles tendon cell density as determined from Achilles tendons sections (4 µm) stained with hematoxylin and eosin (H&E). Data presented as mean±standard error. Representative histology panels are provided for each group.

Cell counts from the H&E stains were not influenced by OVX or genistein (p>0.05). Although an extensive histological evaluation of morphology was not conducted, no gross differences in morphology across groups was noted. Using fibroblasts derived from the Achilles tendon of Intact rats, administration of estradiol increased cell proliferation relative to control (p<0.05) while genistein reduced cell proliferation (p<0.05, Figure 8). The effect of estradiol on cell proliferation was blunted only in the cells treated with the ER-α antagonist MPP (Figure 8). The effect of genistein on cell proliferation was not impaired by either ER-α or ER-β antagonism (Figure 8).

**Figure 8:**
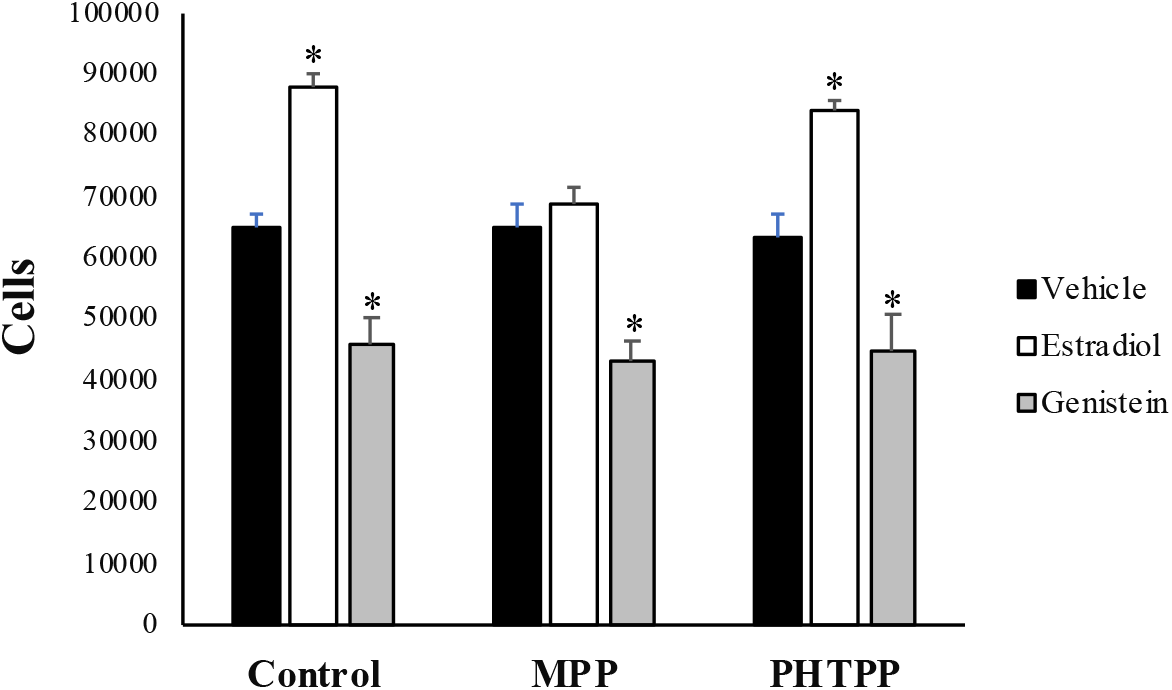
Cell counts following 48 hour treatment. Data presented as mean±standard error. *p<0.05 vs. vehicle.

## DISCUSSION

In our previous work, we demonstrated that loss of estrogen due to OVX leads to a large decline in rat Achilles tendon collagen, an effect that is prevented by administration of genistein (Ramos et al. 2012). In the current investigation, we follow-up on these findings by determining the impact of OVX and genistein treatment on tendon functional properties, while also exploring the effects of OVX and genistein on tendon ECM regulators and cell proliferation. In line with our Achilles tendon collagen findings (Ramos et al. 2012), tendon fascicle failure stress tended to be lower after OVX and genistein treatment led to higher failure stress. Tendon fascicle modulus and energy density were also modestly higher in OVX genistein treated rats. Surprisingly, even with the large loss of Achilles tendon collagen noted in our previous work (Ramos et al. 2012), we did not observe an effect of OVX or genistein on Achilles tendon Col1a1 or Col3a1 expression. Further, key genes associated with ECM degradation were not altered by OVX or genistein treatment. We did, however, observe a modest increase in Dcn expression, a proteoglycan involved with regulating collagen fibril formation and inhibiting cellular proliferation (Yoon and Halper 2005), with genistein treatment. Although no effect of OVX or genistein on the expression of Wnt pathway associated genes was noted, we did observe significant effects of OVX and genistein on Tnmd and Pcna expression, which are key modulators of cellular proliferation (Dex et al. 2016). Interestingly, neither OVX nor genistein altered cell counts when evaluated by H&E staining but genistein did suppress cell proliferation *in vitro*. Further, the effect of genistein on cell proliferation *in vitro* did not appear to require estrogen receptor signaling. Based on our gene data, we hypothesize that the effects of OVX and genistein on tendon collagen may be driven more by changes in cellular proliferation genes rather than direct effects on expression of ECM-related genes. Future studies with more specific measures of cellular proliferation and enzyme activity are needed to further investigate this hypothesis.

Collagen is the primary protein constituent of tendon and correlates strongly with tissue mechanical properties (Ducomps et al. 2003). Therefore, it seemed likely that tendon fascicle mechanical properties would be impaired in OVX rats, given our previous report demonstrating a large loss of tendon collagen with OVX (Ramos et al. 2012). Consistent with limited previous studies (Circi et al. 2009), tendon fascicle failure stress tended to be lower after OVX. Genistein treatment also resulted in higher modulus and energy density when compared to untreated OVX rats. However, we did not observe effects on several other critical functional variables such as stiffness. Changes in tendon stiffness can alter musculoskeletal performance during functional activities (Bojsen-Moller et al. 2005), but our findings suggest that loss of estrogen may not alter tendon function under submaximal loading conditions. Importantly, genistein administration, at amounts easily obtained in the diet, was effective at preserving tendon fascicle failure stress to similar levels as those of intact rats. Our findings provide preclinical evidence that genistein, at doses that can be easily obtained from dietary consumption, may be useful for minimizing the impact of estrogen loss on tendon functional properties. The conclusions of our mechanical findings should be considered with some caution as we are comparing our tail tendon fascicle findings to our previous collagen work in the Achilles tendon (Ramos et al. 2012).

A second objective of the current investigation was to further develop our understanding of the mechanisms mediating the effects of estrogen loss and genistein on tendon. Given the large decline in total tendon collagen noted previously (Ramos et al. 2012), we first examined the impact of OVX and genistein on Col1a1 and Col3a1 expression as well as expression of the proteoglycans decorin and betaglycan (Tgfbr3). Surprisingly, we did not observe an effect of OVX or genistein on the expression of collagen genes. This finding suggests that changes in collagen content due to OVX or genistein may be mediated by post-translational modifications of collagen synthesis or breakdown rather than alteration of gene transcription. Specific studies directly assessing collagen synthesis and related signaling pathways would provide further insight.

Although betaglycan expression was not altered by either OVX or genistein, Dcn expression was higher in OVX-Genistein treated rats relative to OVX-Vehicle. Dcn is the most abundant proteogylcan in tendon and a highly expressed transcript (Table 3), with roles modulating fibrillogenesis and mechanical properties (Robinson et al. 2004; Robinson et al. 2005; Yoon and Halper 2005; Dunkman et al. 2013). Consistent with our findings, steroids, such as dexamethasone, have been shown to stimulate decorin expression in fibroblasts without altering betaglycan expression (Kahari et al. 1995). While we are not aware of any previous investigation in tendon demonstrating an ability of genistein to modify decorin transcription or other proteoglycans, genistein has been shown to inhibit versican expression but not decorin and biglycan in arterial smooth muscle cells (Schonherr et al. 1997). Regardless, genistein had only modest effects of proteoglycan expression in our model, and further work is needed to determine if this effect contributes to the noted improvements in tendon function and collagen content.

In addition to collagen and proteoglycans, a large family of MMPs and TIMPs regulate degradation of ECM components including collagen. Previous work by Pereira et al. (Pereira et al. 2010) reported that MMP-2 enzyme activity was greater in the rat Achilles tendon after 3-months of OVX when compared to intact rats. Further, in non-tendon models, genistein has been shown to inhibit induction of MMPs, which degrade collagen and other ECM components (Zhang et al. 2012). The combination of these studies yield the inference that estrogen and genistein may modulate expression of some MMPs. Surprisingly, we did not observe any changes in the expression of several common genes encoding MMP and TIMP enzymes. Comparisons to the current study are difficult due to differences in rat age and strain. Further, we only evaluated tendon gene expression, limiting our comparisons to Pereira et al. (Pereira et al. 2010). As with our collagen interpretation, it is possible that the effects of OVX on MMPs could be post-translational, thus further studies are needed to carefully examine genistein’s impact on ECM enzyme activity.

We also determined the impact of OVX and genistein treatment on cell proliferation markers and expression of genes related to the β-catenin/Wnt signaling pathway. Tnmd and Scx are well-established tendon-specific genes that have several roles in the modulation of fibrilogenesis and cell proliferation (Liu et al. 2014; Dex et al. 2017). Additionally, Pcna is common marker of cell proliferation. Although we did not observe a significant effect of OVX or genistein on total cell counts or Scx expression, Tnmd expression was greater in OVX-vehicle rats when compared to Intact-Vehicle. Genistein treatment resulted in lower Tnmd expression relative to OVX-Vehicle, but Tnmd expression under OVX-Genistein was still greater than Intact-Vehicle. Pnca expression was lower in OVX-Vehicle rats when compared to Intact-Vehicle, but its expression was similar when comparing Intact-Vehicle and OVX-Genistein treated rats. The β-catenin/Wnt signaling pathway has not been extensively examined in tendon but has been shown to modulate Tnmd and Scx in tendon-derived cells (Lui et al. 2013; Kishimoto et al. 2017). Activation of Wnt/β-catenin signaling in tendon-derived cells lead to suppressed expression of Scx and Tnmd (Kishimoto et al. 2017). However, we found no effect of estrogen loss or genistein treatment on expression of Wnt/β-catenin-related genes. Further, in our model, Tnmd and Scx responded differently to OVX and genistein, suggesting that loss of estrogen may not modulate Wnt/β-catenin signaling in tendon.

Interestingly, we noted an increase in Esr1 expression after OVX, an effect that was not normalized by genistein treatment. Work in other tissues observed an increase in Esr1 expression after OVX, albeit tissue specific (Mohamed and Abdel-Rahman 2000). Furthermore, using validated Esr2 probes (Biorad), we were not able to detect expression of Esr2 transcripts in the Achilles. We have also noted no expression of Esr2 in the Achilles tendon using RNA sequencing (unpublished observation). In contrast, others have reported the presence of Esr2 protein in tendon-derived cells with greater expression after OVX (Hsieh et al. 2018). Our ddPCR findings suggest that the effects of estrogen on tendon may rely on Esr1 rather than Esr2. This conclusion was confirmed in our *in vitro* model where estradiol treatment increased cell proliferation but not when cells were concurrently treated with an ER-α (Esr1) antagonist. ER-β antagonism did not prevent estradiol induction of cell proliferation, which is consistent with previous *in vitro* findings (Maman et al. 2016). Additionally, genistein reduced cell proliferation *in vitro* but, surprisingly this effect did not seem to require activation of ER-α or ER-β. Genistein is an estrogen receptor agonist and has a much greater affinity for ER-β (Kuiper et al. 1998). Our data suggest that tendon may lack ER-β (Esr2) and that genistein may modulate tendon collagen and ECM gene expression via non-estrogenic pathways.

Of note, ddPCR technology allows for absolute quantification of mRNA transcripts with a detection limit of a single transcript. Although the relationship between overall transcript expression in a tissue and the importance of a gene to tissue cellular function is not clear, our ddPCR data (Table 3) does provide an interesting observation regarding tenocyte “commitment” to generating transcripts for a specific gene. As might be expected, the number of collagen and decorin transcripts were considerably higher than the other genes evaluated. Interestingly, Intact rats, genes such as Tnmd and Scx were expressed at a level of ~50-250 fold less than Col1a1. Expression of Mmp9, an important modulator of ECM breakdown, that has been examined in several tendon related studies, was surprisingly very low.

In conclusion, we demonstrate that OVX lead to a decline in stress at tendon fascicle failure but other important functional variables (e.g. stiffness) were not impacted. We can also conclude that genistein can reverse the effects of OVX on tendon fascicle failure stress. Even though we previously observed a large reduction in tendon collagen content after OVX (Ramos et al. 2012), our current findings infer that these changes may, in part, be modulation of Tnmd and Pcna rather than collagen genes or expression of ECM degrading proteins. Further, our findings suggest that genistein, although shown to have an affinity for estrogen receptors, does not completely reverse the effects of estrogen loss on the expression of several genes in the rat Achilles tendon and does not require estrogen receptors to impact tendon fibroblast proliferation. Future studies, should examine the impact of combined estrogen and genistein supplementation and included protein measures and evaluation of estrogen receptor activation.

## ACKNOWLEDGMENTS

This publication was funded, in part, by NIH R15 AT00860501A1 to C.C.C., Purdue University Research Initiative Funds to C.C.C, and with support from the Indiana Clinical and Translational Sciences Institute funded, in part, by Award Number UL1TR002529 from the National Institutes of Health, National Center for Advancing Translational Sciences, Clinical and Translational Sciences Award. The content is solely the responsibility of the authors and does not necessarily represent the official views of the National Institutes of Health.

## Declaration of Interest

The authors report not conflict of interest.

## References

Abate M, Schiavone C, Di Carlo L, Salini V. 2014. Prevalence of and risk factors for asymptomatic rotator cuff tears in postmenopausal women. Menopause. 21(3):275–280.

Abdi F, Alimoradi Z, Haqi P, Mahdizad F. 2016. Effects of phytoestrogens on bone mineral density during the menopause transition: a systematic review of randomized, controlled trials. Climacteric: the journal of the International Menopause Society. 19(6):535–545.

Alekel DL, Genschel U, Koehler KJ, Hofmann H, Van Loan MD, Beer BS, Hanson LN, Peterson CT, Kurzer MS. 2014. Soy Isoflavones for Reducing Bone Loss Study: effects of a 3-year trial on hormones, adverse events, and endometrial thickness in postmenopausal women. Menopause.

Anderson JJ, Ambrose WW, Garner SC. 1998. Biphasic effects of genistein on bone tissue in the ovariectomized, lactating rat model. Proc Soc Exp Biol Med. 217(3):345–350.

Bojsen-Moller J, Magnusson SP, Rasmussen LR, Kjaer M, Aagaard P. 2005. Muscle performance during maximal isometric and dynamic contractions is influenced by the stiffness of the tendinous structures [Clinical Trial Research Support, Non-U.S. Gov’t]. J Appl Physiol. 99(3):986–994. eng.

Circi E, Akpinar S, Balcik C, Bacanli D, Guven G, Akgun RC, Tuncay IC. 2009. Biomechanical and histological comparison of the influence of oestrogen deficient state on tendon healing potential in rats. International orthopaedics. 33(5):1461–1466. eng.

Cook JL, Bass SL, Black JE. 2007. Hormone therapy is associated with smaller Achilles tendon diameter in active post-menopausal women. Scandinavian journal of medicine & science in sports. 17(2):128–132. eng.

Council NR. 2011. Guide for the Care and Use of Laboratory Animals. 8th ed. Washington, DC: The National Academies Press.

Dex S, Alberton P, Willkomm L, Sollradl T, Bago S, Milz S, Shakibaei M, Ignatius A, Bloch W, Clausen-Schaumann H et al. 2017. Tenomodulin is Required for Tendon Endurance Running and Collagen I Fibril Adaptation to Mechanical Load. EBioMedicine. 20:240–254.

Dex S, Lin D, Shukunami C, Docheva D. 2016. Tenogenic modulating insider factor: Systematic assessment on the functions of tenomodulin gene. Gene. 587(1):1–17.

Ducomps C, Mauriege P, Darche B, Combes S, Lebas F, Doutreloux JP. 2003. Effects of jump training on passive mechanical stress and stiffness in rabbit skeletal muscle: role of collagen. Acta physiologica Scandinavica. 178(3):215–224.

Dunkman AA, Buckley MR, Mienaltowski MJ, Adams SM, Thomas SJ, Satchell L, Kumar A, Pathmanathan L, Beason DP, Iozzo RV et al. 2013. Decorin expression is important for age-related changes in tendon structure and mechanical properties. Matrix biology: journal of the International Society for Matrix Biology. 32(1):3–13.

Finni T, Kovanen V, Ronkainen PH, Pollanen E, Bashford GR, Kaprio J, Alen M, Kujala UM, Sipila S. 2009. Combination of hormone replacement therapy and high physical activity is associated with differences in Achilles tendon size in monozygotic female twin pairs [Research Support, Non-U.S. Gov’t Twin Study]. J Appl Physiol. 106(4):1332–1337. eng.

Frizziero A, Vittadini F, Gasparre G, Masiero S. 2014. Impact of oestrogen deficiency and aging on tendon: concise review. Muscles, ligaments and tendons journal. 4(3):324–328.

Hansen M, Kongsgaard M, Holm L, Skovgaard D, Magnusson SP, Qvortrup K, Larsen JO, Aagaard P, Dahl M, Serup A et al. 2009. Effect of estrogen on tendon collagen synthesis, tendon structural characteristics, and biomechanical properties in postmenopausal women. J Appl Physiol. 106(4):1385–1393. eng.

Hansen M, Koskinen SO, Petersen SG, Doessing S, Frystyk J, Flyvbjerg A, Westh E, Magnusson SP, Kjaer M, Langberg H. 2008. Ethinyl oestradiol administration in women suppresses synthesis of collagen in tendon in response to exercise [Research Support, Non-U.S. Gov’t]. The Journal of physiology. 586(Pt 12):3005–3016. eng.

Hansen M, Miller BF, Holm L, Doessing S, Petersen SG, Skovgaard D, Frystyk J, Flyvbjerg A, Koskinen S, Pingel J et al. 2009. Effect of administration of oral contraceptives in vivo on collagen synthesis in tendon and muscle connective tissue in young women [Research Support, Non-U.S. Gov’t]. J Appl Physiol. 106(4):1435–1443. eng.

Holmes GB, Lin J. 2006. Etiologic factors associated with symptomatic achilles tendinopathy [Comparative Study Evaluation Studies]. Foot & ankle international / American Orthopaedic Foot and Ankle Society [and] Swiss Foot and Ankle Society. 27(11):952–959. eng.

Hopkins C, Fu SC, Chua E, Hu X, Rolf C, Mattila VM, Qin L, Yung PS, Chan KM. 2016. Critical review on the socio-economic impact of tendinopathy. Asia Pac J Sports Med Arthrosc Rehabil Technol. 4:9–20.

Hsieh JL, Jou IM, Wu CL, Wu PT, Shiau AL, Chong HE, Lo YT, Shen PC, Chen SY. 2018. Estrogen and mechanical loading-related regulation of estrogen receptor-beta and apoptosis in tendinopathy. PloS one. 13(10):e0204603.

Kahari VM, Hakkinen L, Westermarck J, Larjava H. 1995. Differential regulation of decorin and biglycan gene expression by dexamethasone and retinoic acid in cultured human skin fibroblasts. J Invest Dermatol. 104(4):503–508.

Kishimoto Y, Ohkawara B, Sakai T, Ito M, Masuda A, Ishiguro N, Shukunami C, Docheva D, Ohno K. 2017. Wnt/beta-catenin signaling suppresses expressions of Scx, Mkx, and Tnmd in tendon-derived cells. PloS one. 12(7):e0182051.

Kuiper GG, Lemmen JG, Carlsson B, Corton JC, Safe SH, van der Saag PT, van der Burg B, Gustafsson JA. 1998. Interaction of estrogenic chemicals and phytoestrogens with estrogen receptor beta [Research Support, Non-U.S. Gov’t]. Endocrinology. 139(10):4252–4263. eng.

Liu H, Zhu S, Zhang C, Lu P, Hu J, Yin Z, Ma Y, Chen X, OuYang H. 2014. Crucial transcription factors in tendon development and differentiation: their potential for tendon regeneration. Cell Tissue Res. 356(2):287–298.

Lui PP, Lee YW, Wong YM, Zhang X, Dai K, Rolf CG. 2013. Expression of Wnt pathway mediators in metaplasic tissue in animal model and clinical samples of tendinopathy. Rheumatology (Oxford). 52(9):1609–1618.

Maman E, Somjen D, Maman E, Katzburg S, Sharfman ZT, Stern N, Dolkart O. 2016. The response of cells derived from the supraspinatus tendon to estrogen and calciotropic hormone stimulations: in vitro study. Connective tissue research. 57(2):124–130.

Marini H, Minutoli L, Polito F, Bitto A, Altavilla D, Atteritano M, Gaudio A, Mazzaferro S, Frisina A, Frisina N et al. 2007. Effects of the phytoestrogen genistein on bone metabolism in osteopenic postmenopausal women: a randomized trial [Multicenter Study Randomized Controlled Trial Research Support, Non-U.S. Gov’t]. Annals of internal medicine. 146(12):839–847. eng.

Mohamed MK, Abdel-Rahman AA. 2000. Effect of long-term ovariectomy and estrogen replacement on the expression of estrogen receptor gene in female rats [Research Support, U.S. Gov’t, P.H.S.]. European journal of endocrinology / European Federation of Endocrine Societies. 142(3):307–314. eng.

Nedergaard A, Henriksen K, Karsdal MA, Christiansen C. 2013. Menopause, estrogens and frailty. Gynecological endocrinology: the official journal of the International Society of Gynecological Endocrinology. 29(5):418–423.

Patel SH, Sabbaghi A, Carroll CC. 2018. Streptozotocin-induced diabetes alters transcription of multiple genes necessary for extracellular matrix remodeling in rat patellar tendon. Connective tissue research.1–11.

Pereira GB, Prestes J, Leite RD, Magosso RF, Peixoto FS, Marqueti Rde C, Shiguemoto GE, Selistre-de-Araujo HS, Baldissera V, Perez SE. 2010. Effects of ovariectomy and resistance training on MMP-2 activity in rat calcaneal tendon [Research Support, Non-U.S. Gov’t]. Connective tissue research. 51(6):459–466. eng.

Qi S, Zheng H. 2017. Combined Effects of Phytoestrogen Genistein and Silicon on Ovariectomy-Induced Bone Loss in Rat. Biol Trace Elem Res. 177(2):281–287.

Ramos JE, Al-Nakkash L, Peterson A, Gump BS, Janjulia T, Moore MS, Broderick TL, Carroll CC. 2012. The soy isoflavone genistein inhibits the reduction in Achilles tendon collagen content induced by ovariectomy in rats [Research Support, N.I.H., Intramural Research Support, Non-U.S. Gov’t]. Scandinavian journal of medicine & science in sports. 22(5):e108–114. eng.

Robinson PS, Huang TF, Kazam E, Iozzo RV, Birk DE, Soslowsky LJ. 2005. Influence of decorin and biglycan on mechanical properties of multiple tendons in knockout mice. J Biomech Eng. 127(1):181–185.

Robinson PS, Lin TW, Reynolds PR, Derwin KA, Iozzo RV, Soslowsky LJ. 2004. Strain-rate sensitive mechanical properties of tendon fascicles from mice with genetically engineered alterations in collagen and decorin. J Biomech Eng. 126(2):252–257.

Schonherr E, Kinsella MG, Wight TN. 1997. Genistein selectively inhibits platelet-derived growth factor-stimulated versican biosynthesis in monkey arterial smooth muscle cells. Archives of biochemistry and biophysics. 339(2):353–361.

mServices USDoHaH. 2005. Guidance for Industry: Estimateing the Maximum Safe Starting Dose in Initial Clinical Trails for Therapeutics in Adult Healthy Volunteers. In: Food and Drug Adminstration CfDEaR, editor.

Squadrito F, Altavilla D, Crisafulli A, Saitta A, Cucinotta D, Morabito N, D’Anna R, Corrado F, Ruggeri P, Frisina N et al. 2003. Effect of genistein on endothelial function in postmenopausal women: a randomized, double-blind, controlled study. The American journal of medicine. 114(6):470–476.

Tham DM, Gardner CD, Haskell WL. 1998. Clinical review 97: Potential health benefits of dietary phytoestrogens: a review of the clinical, epidemiological, and mechanistic evidence. The Journal of clinical endocrinology and metabolism. 83(7):2223–2235.

Torricelli P, Veronesi F, Pagani S, Maffulli N, Masiero S, Frizziero A, Fini M. 2013. In vitro tenocyte metabolism in aging and oestrogen deficiency [Research Support, Non-U.S. Gov’t]. Age (Dordr). 35(6):2125–2136.

Trock BJ, Hilakivi-Clarke L, Clarke R. 2006. Meta-analysis of soy intake and breast cancer risk. Journal of the National Cancer Institute. 98(7):459–471.

Verheus M, van Gils CH, Keinan-Boker L, Grace PB, Bingham SA, Peeters PH. 2007. Plasma phytoestrogens and subsequent breast cancer risk. Journal of clinical oncology: official journal of the American Society of Clinical Oncology. 25(6):648–655.

Volper BD, Huynh RT, Arthur KA, Noone J, Gordon BD, Zacherle EW, Munoz E, Sorensen MA, Svensson RB, Broderick TL et al. 2015. Influence of acute and chronic streptozotocin-induced diabetes on the rat tendon extracellular matrix and mechanical properties. American journal of physiology. 309(9):R1135–1143.

Yoon JH, Halper J. 2005. Tendon proteoglycans: biochemistry and function. Journal of musculoskeletal & neuronal interactions. 5(1):22–34.

Zhang Y, Dong J, He P, Li W, Zhang Q, Li N, Sun T. 2012. Genistein inhibit cytokines or growth factor-induced proliferation and transformation phenotype in fibroblast-like synoviocytes of rheumatoid arthritis. Inflammation. 35(1):377–387. Eng.

